# User-driven development and evaluation of an agentic framework for analysis of large pathway diagrams

**DOI:** 10.64898/2026.03.10.710813

**Authors:** Marie Corradi, Ivo Djidrovski, Luiz Ladeira, Bernard Staumont, Anouk Verhoeven, Julen Sanz Serrano, Adrien Rougny, Ahmad Vaez, Ahmed Hemedan, Alexander Mazein, Anna Niarakis, Arthur de Carvalho e Silva, Charles Auffray, Egon Wilighagen, Eliska Kuchovska, Falk Schreiber, Irina Balaur, Laurence Calzone, Lisa Matthews, Lorenzo Veschini, Marc E Gillespie, Martina Kutmon, Matthias König, Matti van Welzen, Noriko Hiroi, Oxana Lopata, Pierre Klemmer, Rupert Overall, Tim Hofer, Venkata Satagopam, Reinhard Schneider, Marc Teunis, Liesbet Geris, Marek Ostaszewski

**Affiliations:** Innovative Testing in Life Sciences and Chemistry, HU – University of Applied Sciences Utrecht; Institute of Risk Assessment Sciences (IRAS), Utrecht University; Biomechanics Research Unit, GIGA Institute, University of Liège; Department of Pharmaceutical and Pharmacological Sciences, Vrije Universiteit Brussel (VUB), Vrije Universiteit Brussel; Luxembourg Centre for Systems Biomedicine, University of Luxembourg; Unit of Genetic Epidemiology & Bioinformatics, Department of Epidemiology, University Medical Center Groningen, University of Groningen; Department of Bioinformatics, Isfahan University of Medical Sciences, Isfahan, Iran; Medical Image and Signal Processing Research Center, Isfahan University of Medical Sciences, Isfahan, Iran; Transversal Translational Medicine, Luxembourg Institute of Health; Department of Cancer Research, Luxembourg Institute of Health; Department of Biology, Faculty of Science and Engineering, University of Toulouse, CBI, CNRS; School of Biosciences, University of Birmingham; European Institute for Systems Biology and Medicine; Translational Genomics, NUTRIM, FHML, Maastricht University; IUF – Leibniz Research Institute for Environmental Medicine; Department of Computer and Information Science, University of Konstanz; Luxembourg National Data Service; Computational Systems Biology of Cancer, Institut Curie, U1331 INSERM, Mines ParisTech, PSL; Biochemistry and Molecular Pharmacology, NYU Grossman School of Medicine; Biocomplexity Institute, Department of Intelligent Systems Engineering, Indiana University at Bloomington; Department of Pharmaceutical Sciences, St. John’s University; Maastricht Centre for Systems Biology and Bioinformatics (MaCSBio), Maastricht University; Institute of Biology, Humboldt University of Berlin; Institute of Structural Mechanics and Dynamics in Aerospace Engineering, University of Stuttgart; Department of Systems Biology and Bioinformatics, University of Rostock; Department of Electrical and Electronic Engineering, Kanagawa Institute of Technology; Department of Chemical Toxicology, Norwegian Institute of Public Health

## Abstract

As biomedical knowledge keeps growing, resources storing available information multiply and grow in size and complexity. Such resources can be in the format of molecular interaction maps, which represent cellular and molecular processes under normal or pathological conditions. However, these maps can be complex and hard to navigate, especially to novice users. Large Language Models (LLMs), particularly in the form of agentic frameworks, have emerged as a promising technology to support this exploration. In this article, we describe a user-driven process of prototyping, development, and user testing of Llemy, an LLM-based system for exploring these molecular interaction maps. By involving domain experts from the very first prototyping in the form of a hackathon and collecting both fine-grained and general feedback on more refined versions, we were able to evaluate the perceived utility and quality of the developed system, in particular for summarising maps and pathways, as well as prioritise the development of future features. We recommend continued user-driven development and benchmarking to keep the community engaged. This will also facilitate the transition towards open-weight LLMs to support the needs of the open research environment in an ever-changing technology landscape.

## Introduction

The ecosystem of biomedical knowledge repositories continues to grow in size and complexity. Structured information available in databases of pathway diagrams, knowledge graphs, or interaction databases can support hypothesis generation, inform experimental design, and facilitate interpretation of omics data (I. Mazein et al., 2024). However, navigating resources with different formats, interfaces, and granularities is a challenging and time-consuming task. Additionally, the sheer volume of these knowledge collections poses a challenge for navigating their individual components and relationships.

One of these resources is comprehensive graphical representations of cellular and molecular processes associated with specific organ functions under normal or pathological conditions (A. Mazein et al., 2018; Staumont et al., 2025). Such maps are manually curated, provide information about source literature used for their construction, and follow the standards of Systems Biology Graphical Notation (SBGN) (Le Novère et al., 2009) and Systems Biology Markup Language (SBML) (Keating et al., 2020). They are hosted using the MINERVA Platform (Hoksza et al., 2019), which provides functionalities for visual exploration and programmatic access. Currently a number of such disease and physiological maps are available, describing pathways of neurodegeneration (Fujita et al., 2014), chronic immune and allergic diseases (Acencio et al., 2023, 2026; Lopata et al., 2025; A. Mazein et al., 2021), mechanisms of inflammation and viral infection (Ostaszewski et al., 2021; Serhan et al., 2020), and physiology of liver metabolism (Ladeira et al., 2025) or adverse outcome pathways (A. Mazein et al., 2025). Importantly, repositories hosting molecular interaction maps are distributed and differ in scope and granularity, and can benefit from unified access interfaces to reduce barriers to access and interpretation. This will facilitate visualisation, analysis and modelling of complex molecular mechanisms represented by these maps (Hemedan et al., 2024; Hoch et al., 2022, 2023; Zerrouk et al., 2022, 2024).

Large Language Models (LLMs) have emerged as a promising technology for summarising, analysing, and explaining such structured knowledge. Their applications range from general-purpose foundational models to specialised tasks like building and exploring knowledge graphs or data analysis and hypothesis generation. Direct retrieval of knowledge from LLMs has been evaluated in, e.g., assisting pathway enrichment analysis (Azam et al., 2024), retrieving ageing-relevant knowledge under different hyperparameter configurations (Shakhbazau, 2025), or searching for drug combinations in Alzheimer’s disease (E. J. Wang et al., 2026). For processing structured knowledge, LLM-based workflows have been proposed to generate knowledge graphs from the literature and to process them for prioritising experimental results (Feng et al., 2025) and exploring clinically valid hypotheses (Khan et al., 2025). In parallel, frameworks have been proposed to convert biological databases into structured knowledge graphs (Lobentanzer et al., 2023) and to access information via chatbot interfaces (Lobentanzer et al., 2025). In parallel to specific use cases, LLM-supported architectures were proposed, assisting with access to bioinformatic databases and summary of results (Sokolova et al., 2025), supporting collaborative construction of scientific hypotheses (Jiang et al., 2026), or comprehensive analysis of omics data (Nagarajan et al., 2025). Finally, LLMs were evaluated for their support in curation and annotation of bioinformatic knowledge (Devkota et al., 2025), including pathway databases (Wu et al., 2025). However, the need for a dedicated solution addressing molecular interaction maps is still missing. As these maps are highly interactive resources with a range of different applications, an LLM-based system should be developed and evaluated by directly engaging the target user group.

In this article, we describe a user-driven process of prototyping, development, and user testing of Llemy, an LLM-based system for exploring large pathway diagrams. Llemy is an online system built for exploratory analysis of user-built repositories of molecular interaction maps. We evaluated Llemy’s usability and its perceived performance by having a group of users conduct an exploratory analysis of various molecular interaction maps. We gathered information on the system’s performance at the level of individual user prompts and collected overall user feedback via a survey completed after their work. Based on these results, we identify key tasks performed by the users, and assess their perceived performance. We demonstrate a need for a robust framework for open-ended evaluation of tasks involving interactive diagram exploration and analysis. The results of this evaluation provide insight into the design of future LLM-based systems for exploring and analysing molecular interaction maps and indicate the types of tasks on which such systems may underperform.

This article is structured as follows. In the Methods section, we introduce the hackathon in which Llemy was prototyped, the user-driven approach to its development, its architecture, and the evaluation process. In the Results section, we briefly describe the user interface and analyse user feedback data. Finally, we discuss the results and provide an outlook for future work.

Llemy can be accessed via https://llemy.vhp4safety.nl, and its code repository is available at https://github.com/ontox-project/Llemy (see Data availability).

## Methods

Llemy was developed in several steps, following a user-centric design. First, a prototype was developed during a two-day hackathon. This prototype was then refined and evaluated by a group of users involved in the development and analysis of molecular interaction maps.

### Prototype development

The project was initiated during a two-day hackathon held at the University of Liège, Belgium, on May 8-9, 2025. The aim was to design a system that facilitates exploration of molecular interaction maps, with liver lipid and bile metabolism (Ladeira et al., 2025) as use cases. These maps were developed as a part of the ONTOX project (Vinken et al., 2021). The hackathon group involved hepatotoxicologists, curators of the maps, computational biologists, and LLM experts.

User requirements and the system architecture were established based on user prompts prepared by the domain experts. These prompts were then used in iterative prototyping and evaluation. The prompts included i) queries with scope beyond the contents of the map, and ii) queries containing false premises, designed to evaluate the ability to identify and address incorrect assumptions (see Supplementary H1).

The final hackathon prototype consisted of two separate agents collecting information in parallel: one from the map hosted on the MINERVA Platform, and one from Perplexity using deep research (accessed on 08.05.2025). Then, a third synthesis agent (OpenAI GPT4.1) combined and summarised the findings. Commercial LLMs were chosen for quick prototyping. The whole prototype was containerised in a Docker application with the possibility to deploy it locally and provide API keys for both LLMs. This architecture was inspired by the O-QT assistant (Djidrovski et al., 2025).

### System architecture

Llemy was built entirely in Python, using GPT4.1-nano, Streamlit (v1.50.0) for the front-end, and LangChain (v0.3.27) as the agentic backend. Llemy is deployed publicly on a cloud platform (https://llemy.vhp4safety.nl/), and works with user-provided API keys. It is also available as a Docker container for local deployment. In the deployed version, API keys are stored only for the duration of a user session to ensure security. To avoid context rate limits, a follow-up prompt using the previous answer is not possible in the current version as of the publication of this paper. However, the entire chat history from a given session is visible. Llemy combines user prompt with map data and instructions output, focus, and inclusion of literature references. The full instructions are available in the file **`workflow.py`** on the GitHub repository (https://github.com/ontox-project/Llemy).

### User study design

Llemy was evaluated by a group of 25 users recruited via Disease Maps Community channels (https://disease-maps.io/). The users worked with the system by performing queries to a selected pathway or disease map, using either a pre-constructed set of prompts (see Supplementary U1), or prompts provided by the users. Users were provided with API keys and asked to evaluate the system’s performance after each prompt. Performance metrics were accuracy, conciseness, and reliability, evaluated on a five-point scale. Users were also able to provide free-text comments on the system’s performance for each metric. The API keys for user testing were provided by ELIXIR Luxembourg Node. At the end of their testing session, the users were asked to fill out a questionnaire summarising their overall assessment of the performance of Llemy (see Supplementary U2).

Two separate datasets were collected for analysis. One consisted of individual prompts, system replies, and user feedback on the accuracy, conciseness, and reliability of the responses (Prompt Dataset). The other dataset contained user replies to the final questionnaire (Summary Dataset). They were treated as separate in downstream analyses.

#### Prompt Dataset analysis

The Prompt Dataset contains 157 entries, including the user prompt, the response of the system, the response time, and the evaluation of the response in terms of accuracy, conciseness, and reliability, rated on a 5-point scale (see Supplementary Table U3). One prompt, intentionally designed to test the system’s ability to remain within the scope of its source maps, was removed from the subsequent analysis. To understand the prompt landscape, 114 unique prompts were categorised into three categories. The categories were established post hoc based on the content of the prompts. Each prompt was assigned a category based on a consensus among three independent experts. Score distributions were summarised, both aggregated across all categories and stratified by prompt category, to examine potential variation in perceived system performance across different categories. Free-text comments provided for each prompt evaluation were grouped by associated metric score and analysed qualitatively. There were no comments for conciseness.

#### Summary Dataset analysis

The Summary Dataset gathers a general evaluation of the system from 19 unique users. The data collection was fully anonymous. Users were asked about their experience using molecular interaction maps and LLMs. They were then asked to estimate how long they spent testing the app, and what type of prompts they entered. Further feedback on ease of use, productivity and overall assessment were collected using both a 5-point scale and free text comments.

#### Statistical analysis

All analysis was performed in R version 4.5.2. To analyse the impact of response time on performance metric, a cumulative link mixed model (ordinal package, clmm method, link = “probit”, nAGQ = 10) was fitted on Accuracy, Conciseness and Reliability metrics (jointly), with a random intercept per user to account for within-user correlation. A cumulative link mixed model was chosen because of high inter-metric correlation: Spearman correlation between Accuracy and Conciseness is 0.53, between Conciseness and Reliability is 0.43, and between Accuracy and Reliability is 0.71.

To examine differences in Accuracy, Conciseness, and Reliability across Categories, post-hoc pairwise comparisons, using FSA package, were conducted using Dunn’s test with Holm correction for multiple comparisons This non-parametric test was applied separately for each dependent variable to identify which specific category pairs differed significantly while controlling for family-wise error rate.

## Results

### Implementation of Llemy

Llemy consists of two parallel processes (Figure 1) following the prompt provided by the user, and the selected map. First, the user prompt is enriched with instructions for the LLM ensuring contextualised output, scientific focus, and inclusion of literature references. Second, a retrieval agent uses the MINERVA API (Hoksza et al., 2019) to get contents of the map selected by the user: elements (nodes), reactions (edges), and annotations. The system provides access to public maps available under a permissive licence shared via a repository of molecular interaction maps, MINERVA Net (https://minerva-net.lcsb.uni.lu/) (Gawron et al., 2023), and a set of predefined prompts. The contents are concatenated into a single text object, including element identifiers, reaction types and identifiers of supporting literature.

**Figure 1:**
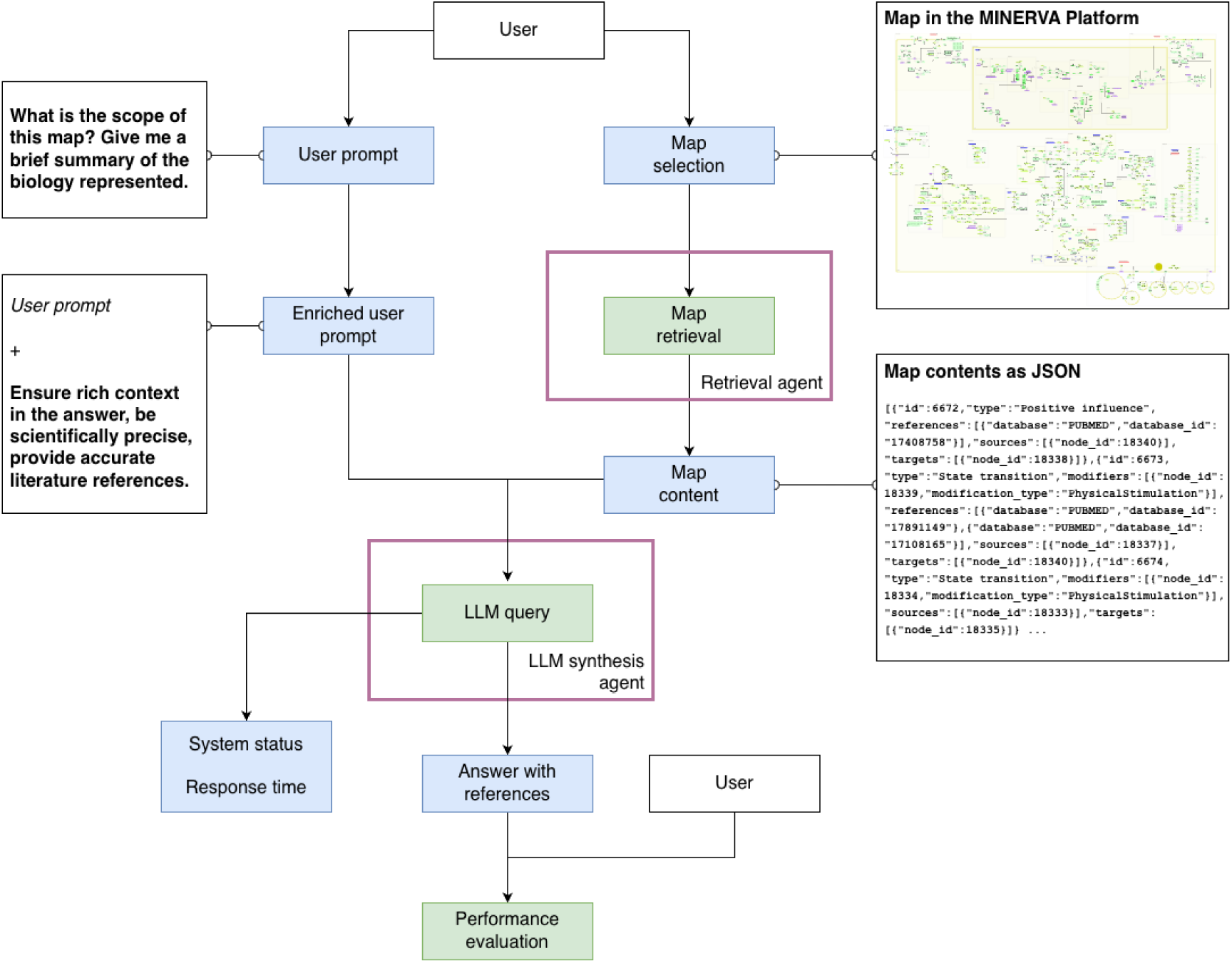
The workflow of Llemy. Users provide their prompt and select a map they want to process. Then, the prompt is enriched with instructions for the content of the answer (see full system prompt in Supplementary P1), and, in parallel, map data is fetched from the corresponding MINERVA Platform. Both enriched prompt and processed map data are combined and used as the LLM prompt. The answer is provided to the user for evaluation, and system performance logs are collected. Green: process, blue: data.

This text object is then given to a “synthesiser agent” as a context, together with the enriched user prompt. The response is formatted post-hoc by adding links to specific parts of the map of interest: when the LLM cites an element or reaction identifier in a given sentence, they are isolated and collated to a clickable link after the end of that sentence. A robust error-handling and logging layer is integrated throughout the workflow: API responses are normalised into explicit states (e.g., success, error, no_data_found, skipped), retrieval errors are passed to the synthesiser as structured context, and both the synthesised response and raw map payload are returned for traceability. This makes each answer auditable and allows users to verify exactly which MINERVA-derived data supported the final output. Response times and carbon emissions for each query were also calculated; for the latter, unfortunately, the calculation mechanism failed in 90% of the cases so we were unable to draw any conclusions.

Llemy is deployed as a service on the VHP4Safety cloud (https://llemy.vhp4safety.nl/). Upon landing on the page (see Figure 2), the user is prompted to enter an OpenAI API key. To provide user feedback, a password is required to ensure privacy, and consent to collect prompt and response data to analyse the performance of the system. If the user previously agreed to the recording of their data, a brief feedback form is displayed to rate the accuracy, conciseness, and reliability of the answer on a 1 to 5 scale, and provide a free text comment on the result of a particular query. Llemy can be used without providing feedback.

**Figure 2:**
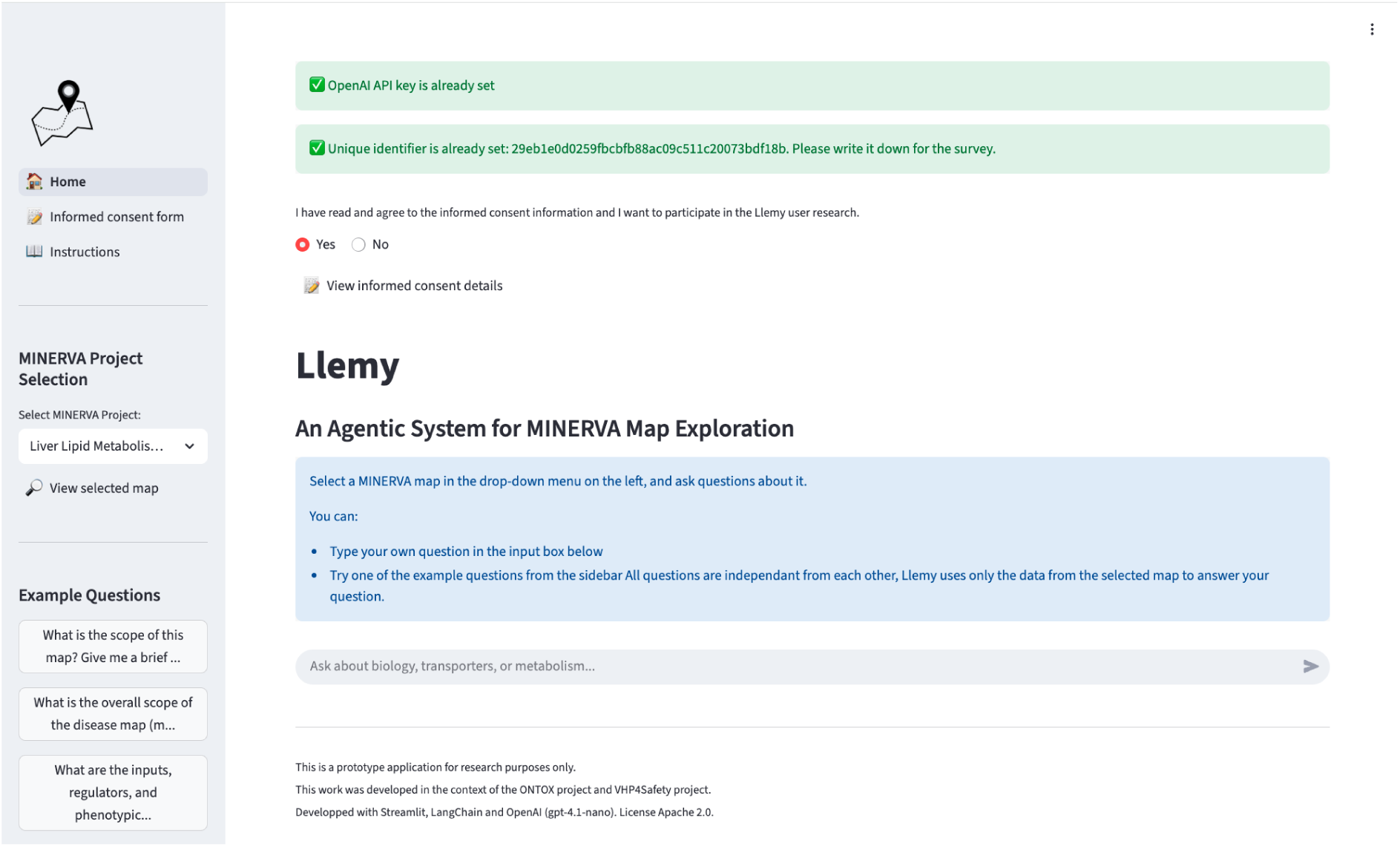
Screenshot of Llemy’s welcome page. The left-side panel comprises i) a section with links to the home page, informed consent form, and user manual; ii) a map selection drop-down menu, and a link to the selected map; iii) a list of example prompts (see Supplementary U1 for a complete list).

### User study

#### User feedback for individual prompts

We assessed the perceived performance of the system across three metrics – accuracy, conciseness, and reliability – based on user feedback (see Figure 3). Medians for these three metrics were 4, 3, and 4, respectively. We performed a cumulative linked mixed model (see Methods) to check if any of these metrics were affected by the response time. Response time had a significant negative effect on ratings (β = −0.34, SE = 0.06, z = −6.06, p < 0.001), indicating that longer response times were associated with lower perceived quality. This effect was uniform across Accuracy, Conciseness and Reliability. On average, users rated Conciseness significantly lower than Accuracy (β = −0.37, p = 0.002), while Reliability did not differ from Accuracy (β = −0.12, p = 0.34).

**Figure 3:**
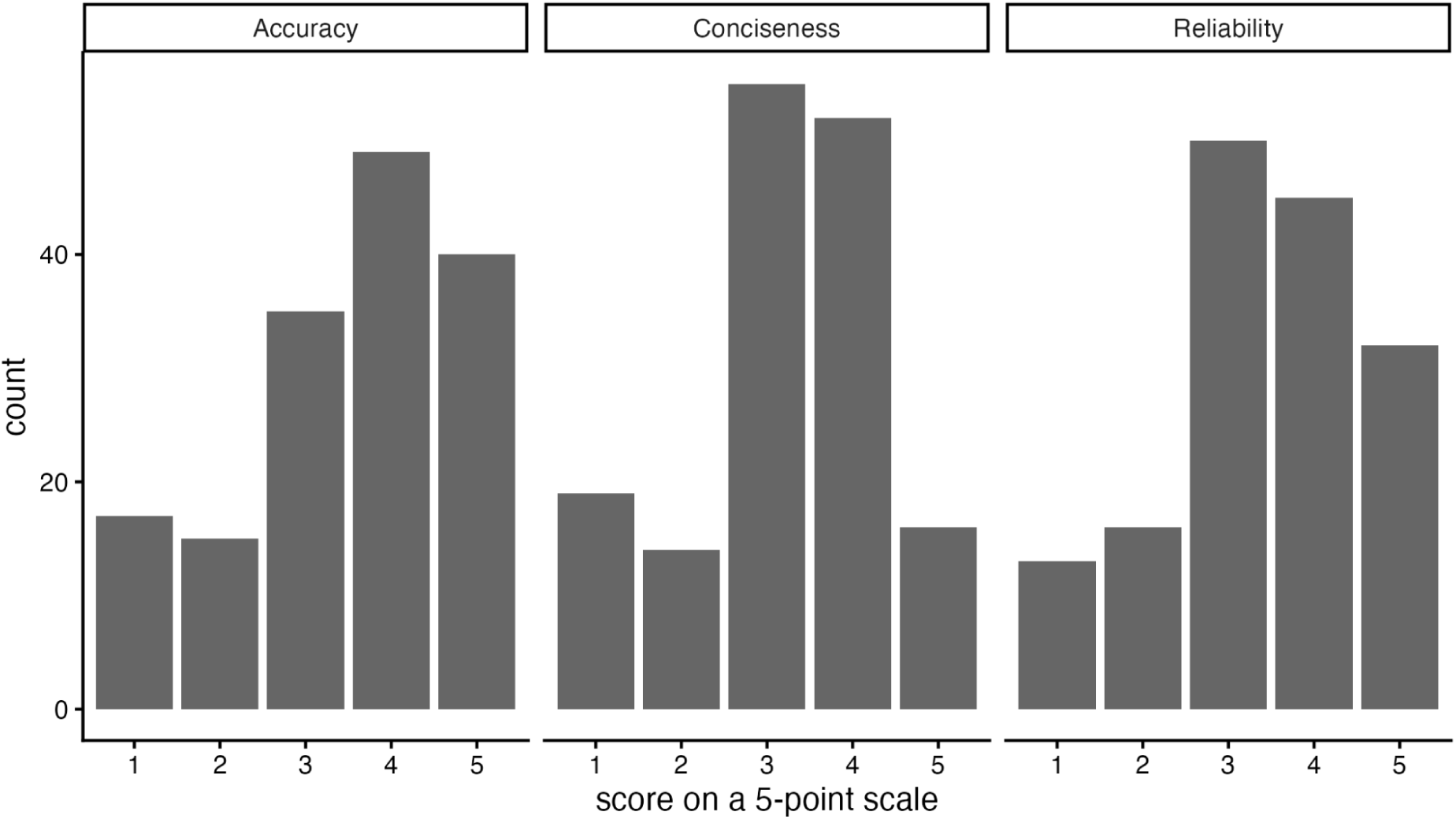
Score distribution by evaluation criterion.

Individual prompts were grouped into three categories (see Methods), as shown in Table 1, together with the number of prompts per category. Stratified analysis of evaluation metrics using post-hoc Dunn test (see Methods), revealed no significant performance differences across categories. Mean accuracy, conciseness, and reliability were the highest for the “Summarise” task (Figure 4), and the “Find” category exhibited a broader distribution of scores, showing lower evaluations for all scores, possibly reflecting the complexity of querying molecular interaction maps with an LLM when the content of the map is in the prompt.

**Figure 4:**
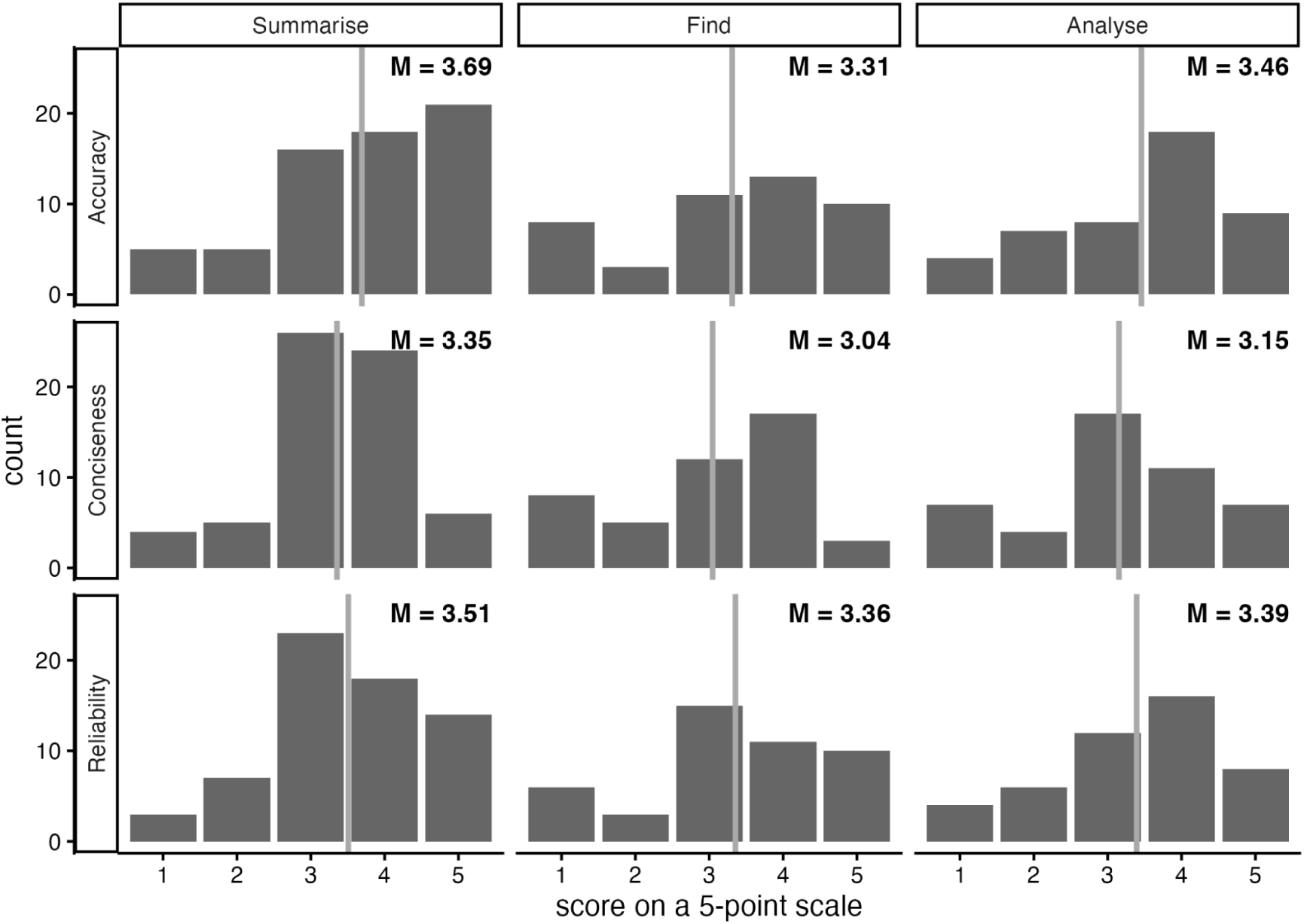
Distribution of tasks performed by the users testing Llemy, grouped into three categories. M = average.

**Table 1:** Categories of prompts in the Prompt Dataset, and their short definitions.

| Prompt category | N | % | Definition |
| --- | --- | --- | --- |
| Summarise | 65 | 41.7 | Summary of a map, mechanisms involving specific pathways, elements or interactions (e.g., autophagy, bile acid metabolism, signalling pathways), or mechanisms missing from the map. |
| Find | 46 | 29.5 | Retrieval of specific map elements, elements having specific properties (e.g., kinases, ligands, enzymes, transcription factors, protein interactors), or upstream or downstream targets (e.g., receptors, organelles, phenotypes) outside of the map. |
| Analyse | 45 | 28.8 | Analysis of the map or its parts (e.g., count specific elements, calculate connectivity of the network) or reason about the consequences of perturbing the molecular network (e.g., drug effects, disease-related perturbations) |

Qualitative analysis of individual user comments revealed distinct patterns associated with score levels. For higher accuracy scores (4-5), users acknowledged the system’s ability to provide comprehensive summaries, correctly identify pathway connections across submaps, and appropriately acknowledge limitations when specific entities were absent from maps. For lower accuracy scores (1-2), users most frequently cited fundamental factual errors, failure to locate existing map content, and fabricated or mismatched map reaction references. Common criticisms included misunderstanding map element types, missing available annotations, and instances where the system failed to recognise entities present under synonyms. There were no individual comments on the conciseness of results.

Reliability scores were strongly influenced by the quality of the reaction hyperlinks to the source content, coming from maps hosted in the MINERVA platform. High reliability scores (4-5) corresponded to systematic and verifiable referencing, allowing systematic verification. Low reliability scores (1-2) were associated with broken or non-resolving links and inconsistent behaviour across repeated queries, notably varying structure and scope of provided references to the source map, from insufficient to unnecessarily repetitive.

Several cross-cutting themes emerged from user feedback. First, synonym and nomenclature handling presented recurring challenges when the system failed to recognise entities represented by HGNC (HUGO Gene Name Committee) names rather than common abbreviations. Second, users valued contextual awareness, noting that responses sometimes missed map-specific or domain-specific framing (e.g., failing to mention organ context for an organ-specific map). Third, users at intermediate score levels (3) noted that answers were accurate but incomplete, and challenging to assess due to a lack of proper references to the source material.

Overall, we summarised individual user comments having particular score levels as listed in Supplementary U4. No individual comments on the conciseness of the system were submitted.

#### Overall user feedback

Results of the user survey showed a varied distribution of users, curators, and developers of molecular interaction maps. The relatively low number of responses in the “user” group can be explained by the fact that most users also had experience as curators or developers. Almost all users had prior experience with LLMs. All users tested the system for more than 15 min, half of them spent more than an hour, and 75% estimated that they saved time by using Llemy. Another important feedback was concerning the variability of the output for similar queries, where on a 1 to 5 scale, all users except one reported variability of 3 and higher.

## Discussion

### Advantages of the LLM-based approach

The potential of LLMs to accelerate research is a topic of intense investigation – including in the field of molecular pathways and their regulation. In this work, we explore the capability of LLM-based systems to process, summarise, analyse, and extend structured, focused knowledge repositories describing molecular pathways involved in human physiology and pathology. We introduce Llemy, a system for summarising knowledge graphs based on user prompts. Our approach resembles proposed LLM solutions to refine knowledge graphs from literature (Feng et al., 2025; Khan et al., 2025), but the sources of the system are manually curated diagrams. This allows Llemy to directly reference diagram elements and interactions, thereby improving the system’s transparency and facilitating exploration of the original content based on the system’s response.

### User-focused analysis

Our approach focused on the user perspective in the design and evaluation of the system. The system was designed and prototyped during a hackathon, when domain experts directly formulated prompts they would ask, and evaluated replies on-the-fly. The finalised system was then tested by an independent group of 25 users at the level of individual prompts, and via a survey of their overall evaluation. Based on 156 individual prompts, we proposed three categories of prompts posed to the system and showed that tasks with higher complexity are asked less frequently and receive less positive feedback. Although no statistical differences were observed, we’ve noticed comparatively higher scores in the “Summarise” category. Based on these categories, Llemy workflows can be tailored to the tasks performed by users. This is similar to the functionality of the Alvessa system proposed in (Sokolova et al., 2025), but focuses on the nature of the task rather than on specific resources to be queried based on the system prompt.

### User perception of an LLM-based system

A summary user survey completed by 19 of 25 participants rated the system as highly accessible; however, the utility rating was more moderate, particularly among map users. This may be explained by the fact that the users less familiar with the content processed by Llemy can give more favourable evaluations to its answers. All users generally rated the usability of Llemy as high (see Figure 5), with over 80% rating it a 4 or 5. It is worthwhile to note that users-only were quite split. Over 70% of users found Llemy useful (score of 4 or more).

**Figure 5:**
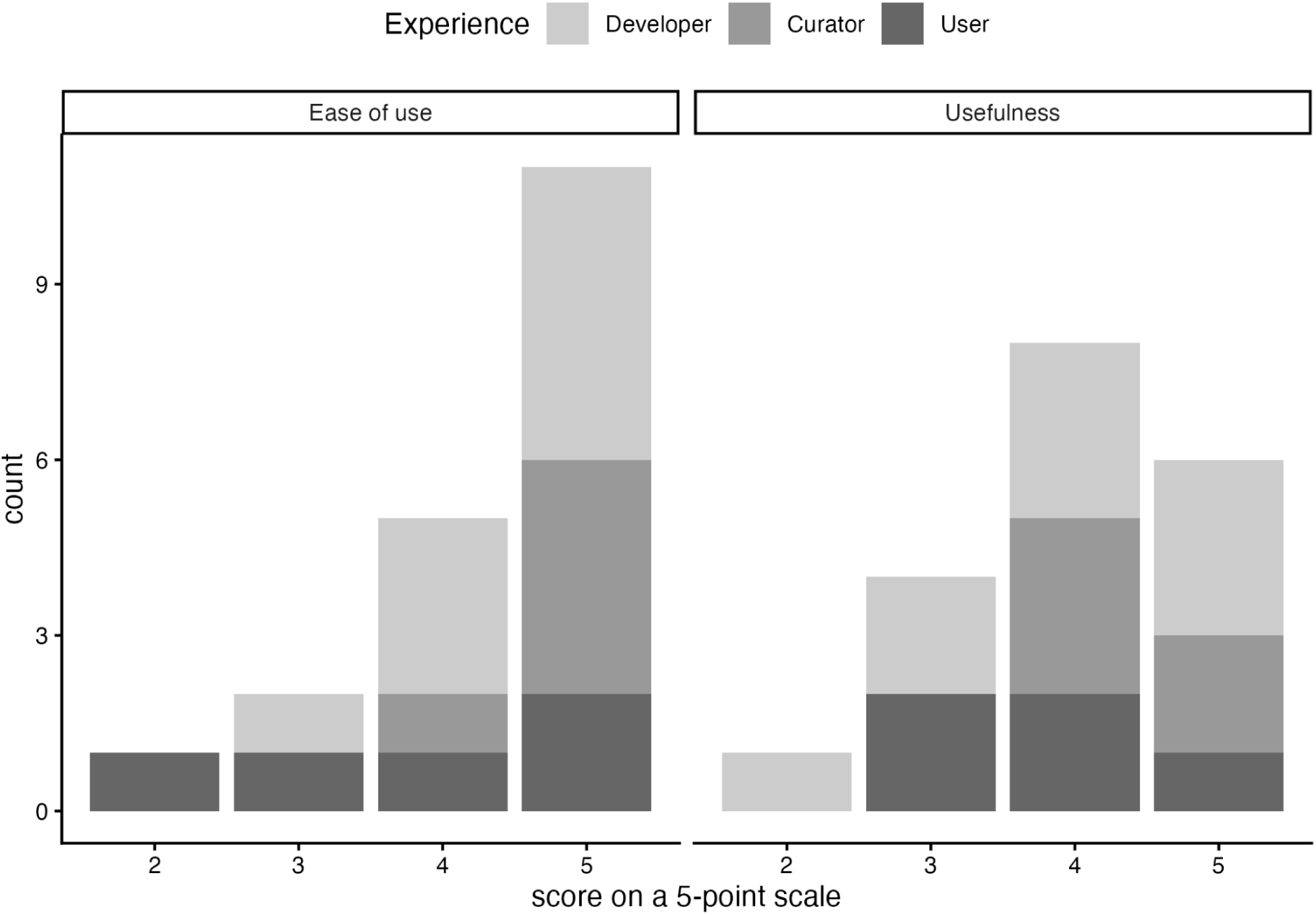
Usefulness and usability based on experience with molecular interaction maps. Developer – participants building workflows including maps; Curator – participants creating maps; User – participants using the maps for knowledge retrieval and data visualisation. In case of multiple roles, the highest level of experience – Developer, then Curator, then User – is taken into account.

Reproducibility of results and the sustainability of the system are key considerations in developing systems such as Llemy. All users except one reported a marked variability of the system output, of at least 3 points on the 1 to 5 scale, in line with other reports (J. J. Wang & Wang, 2025). In order to improve this situation, open-weight models may be considered, contrary to a commercial LLM used currently. Importantly, open-weight LLM models show promising performance when compared to commercial solutions (Stachura et al., 2025), however, use of open LLMs comes with the need for a dedicated infrastructure and associated costs. Moreover, tests of locally deployed, open-access, and open-weight LLMs are necessary to compare the tradeoff between quality and cost of use, which in turn requires benchmark datasets. Our open-ended user survey may inform the design of such a system, similar to foundational model benchmarks (Laurent et al., 2024) used in Sokolova et al. (2025).

### Roadmap for the system and its evaluation

Overall, Llemy and similar systems have the potential to reduce the complexity barrier when working with complex diagrams. Based on the user feedback in both the Summary and Prompt datasets, we drafted a roadmap to support future improvements. We attributed priorities based on the frequency with which users requested the feature. In the short term, systems architecture should be updated to improve response time and accuracy of the provided references. Afterwards, dedicated workflows for summarising, querying, and analysing the map should be introduced, with task granularity matching the user study. Finally, programmatic integration of Llemy functionalities with the GUI of the MINERVA Platform using plugins (Hoksza et al., 2019) and workflows using the Model Context Protocol (Hou et al., 2025; Kuehl et al., 2025) will streamline the interaction of LLMs with the content of maps. This should be supported by use of open LLMs to address the needs of the open-access research environment.

User interaction is an important part of the Llemy project, and it can be improved by acting directly on provided feedback. Llemy needs improvements supporting term disambiguation and feedback for prompts that cannot be answered, as well as better linking of references to original publications or map content used in the answer. Better transparency into requested data and its use in answering user queries will improve understanding and evaluation of the system’s behaviour. These steps will lead to an improved user evaluation system, likely combining benchmarks specific to proposed tasks with an open-ended scoring system.

The user-based evaluation system applied for Llemy requires refinement and further development, together with the changing landscape of technology and its possible applications. Open-ended user testing in computational biology is a limited, but expanding area (Mitchener et al., 2025), which can inform the development of highly interactive and mulitpurpose resources like molecular interaction maps. The framework proposed in this article is limited to three metrics of performance, which nevertheless allowed us to post-hoc identify main tasks explored by the user community. This informed the roadmap of Llemy development, and can be expanded into a set of task-specific evaluations, combining structured and open-ended scenarios of map exploration and analysis. Evaluators from communities of interest should be iteratively engaged when such refinements become available, including pathway and network interaction databases, like Reactome (Ragueneau et al., 2026), WikiPathways (Agrawal et al., 2024) or SIGNOR (Lo Surdo et al., 2026), and synergising with relevant efforts (Mohammadi et al., 2025). The timeline of future user testing depends on system development, but can be supported by an engaged community, tailoring LLM support to user needs.

### Study limitations

Our study faces a number of limitations. First, the limited number of users and the recruitment process may have affected the survey’s overall evaluation and conclusions. This can be seen in the relatively low number of participants in the “user” group, i.e., people with no prior knowledge about development or analysis of molecular interaction maps. Second, Llemy features an inconsistent output, which affects the evaluation by its users. The answers vary for prompts having the same content and structure, including missing references, varying answer size, or overwhelming and unnecessary repetitions. This is a known behaviour of LLM-based systems for complex tasks (J. J. Wang & Wang, 2025), and is beyond the scope of our current work. Finally, due to technical limitations, the size of the content processed by Llemy is restricted by the LLM capabilities, resulting in a limited scope of user queries. A potential mitigation for this issue would be to move from a long-context architecture to a (graph) retrieval-based approach, using solutions akin to those developed by Lobentanzer et al. (2025).

## Conclusions

Llemy shows a promising use case for the usage of LLMs to interact with biological databases, particularly as a summarisation tool for an entry point into molecular interaction maps. The user-driven approach we took is important for evaluating the needs and perception of LLM-based systems in science, and yielded valuable insights on further development, in particular for more advanced tasks. With a rapidly changing landscape of available LLMs, it is important to construct sustainable systems that can be benchmarked when underlying models change. User-driven development and open-ended benchmarking systems can support these LLM comparisons, especially for complex tasks using interactive tools. Future plans also include re-running the study with an extended framework, as well as recruiting more uniformly distributed participants for a more generalisable evaluation and a better coverage of possible tasks.

## Disclosure of Ethical Statements

N/A

## Consent for publication

All authors agree to the publication of this study.

## CRediT author statement

Conceptualization: MC, ID, LL, BS, AV, JSS, MT, LG and MO.

Methodology: MC, ID, LL, BS, AV, JSS, MT, LG and MO.

Software: MC, ID and MO.

Validation: MC, ID, LL, BS, AV, JSS, AR, AV, AH, AM, AN, ACS, CA, EW, EK, FS, IB, LC, LM, LV, MEG, MK, MKönig, MW, NH, OL, PK, RO, TH, MT, LG and MO.

Resources: MC, ID, LL, BS, AV, JSS, VS, RS, MT, LG and MO.

Writing – Original draft: MC, ID, LL, BS, AV, JSS, MT, LG and MO.

Writing: review & editing: MC, ID, LL, BS, AV, JSS, AR, AV, AH, AM, AN, ACS, CA, EW, EK, FS, IB, LC, LM, LV, MEG, MK, MKönig, MW, NH, OL, PK, RO, TH, VS, RS, MT, LG and MO.

Supervision: MT, LG, MO.

Funding acquisition: VS, RS, LG, and MT.

## Acknowledgements

Online browsing is supported by the MINERVA team (https://minerva.uni.lu) at the Bioinformatics Core of the Luxembourg Centre for Systems Biomedicine. The work presented in this article was supported by the ELIXIR Luxembourg tools and services. (https://elixir-luxembourg.org).

## Data availability

Llemy’s code and documentation are available at GitHub (https://github.com/ontox-project/Llemy), under the Apache 2.0 License (https://www.apache.org/licenses/LICENSE-2.0). A public instance is hosted at https://llemy.vhp4safety.nl/.

## List of abbreviations

API: Application Programming Interface

GUI: Graphical User Interface

HGNC: HUGO Gene Name Committee

LLM: Large Language Model;

MCP: Model Context Protocol;

MINERVA: Molecular Interaction NEtwoRk VisuAlization

SBGN: Systems Biology Graphical Notation;

