## Supplementary H1 for "User-driven development and evaluation of an agentic framework for analysis of large pathway diagrams"

### Supplementary H1: Hackathon prompts

List and rationale of example prompts used to test the first Llemy prototype:

- List three specific inhibitors of CD36 and tell me which general molecular processes would be mainly impaired.
    - Rationale: Identification and disambiguation of specific proteins within a map, linking them to external drug databases (e.g., DrugBank), and summarizing complex downstream processes.

- Tell me an alternative to FATP2 if I want to test for the functionality of bile canalicular efflux.
    - Rationale: Identification of unrelated/incorrect input. FATP2 is related to fatty acid uptake and has no known link to bile canalicular efflux.

- What protein or protein combinations should I measure if I want to assess HDL secretion?
    - Rationale: A question that implies various steps: the disambiguation of high-density lipoprotein (HDL), the separation of its basic proteic components, the search of the different techniques to measure each of them (from previous trained knowledge), and the final output of prioritization of these techniques.

- In what organelle are fatty acids mainly stored in a cell system not expressing FABP1?
    - Rationale: Assessing the relationship (and causality) between non-existent fatty acid transport (FABP1), the role of isoforms when a primary transporter is absent, and storage (compartment) as two separate entities in the physiological maps.

- What are the differential molecular processes inhibited when PPAR alpha and PPAR gamma are inhibited?
    - Rationale: Complex question that involves the identification of potential pathways in the physiological maps related to PPAR inhibition, and flagging out related pathways to two nuclear receptors from the same family. This requires the system to cross-reference the PM with external transcription factor and target gene databases (e.g., OmniPath) to compare distinct molecular outcomes.

- Is PPAR alpha involved in the uptake of fatty acids?
    - Rationale: This tests the system's ability to identify both direct and indirect downstream events in the map and potentially incomplete map data.

- How is BSEP related to fatty acid beta oxidation?
    - Rationale: Identification of an a priori non-existing relationship between BSEP and fatty acid beta oxidation.

- How does FATP1 inhibition in the liver affect the homeostasis of cholesterol?

    - Rationale: Capability of the system to discern between potential relationships or far-fetched ones. FATP2 and FATP5 are primarily expressed in the liver, so the contribution of FATP1 (more common in liver cancer cell lines) might be limited in comparison. Moreover, the FATP family is related to the transport of fatty acids, therefore with limited direct relation to cholesterol.

- Which cell types should I include in an in vitro model of steatohepatitis?
    - Rationale: Complex relationship between the complex process of steatohepatitis, roughly involving steatosis and inflammation, and the identification of the cells involved (hepatocytes, stellate cells and Kupffer cells). This question is out of the scope of the liver maps used in the design phase, so it test the system's ability to discriminate what can be answered with the context data from what is out of the map.

- What is the abundance of FATP transporters in the liver?
    - Rationale: This evaluates the system's ability to differentiate between and integrate relative abundance from PM overlay datasets when available, while distinguishing between protein and RNA data. This type of data is not retreived by the MINERVA agent, so the answer is out of the scope of the context data and the answer is expected to aknowledge that.

- Is valproic acid an inhibitor of OTG2?
    - Rationale: Identification of unrelated/incorrect input; OTG2 does not exist as protein.
