## Supplementary P1 for "User-driven development and evaluation of an agentic framework for analysis of large pathway diagrams"

### Supplementary U4: Grouping of user’s comments by score ranking for Accuracy and Reliability

Accuracy:

- Score 1: Fundamental factual errors (incorrect statements about map contents, failure to locate existing entities, fabricated references);
- Score 2: Partially correct with significant errors (initial elements accurate but subsequent content flawed, failure to recognise map structure, fabricated supporting text);
- Score 3: Generally reasonable but incomplete (correct directionality with gaps in coverage, missed existing content under alternative nomenclature, difficult to verify without references);
- Score 4: Accurate with minor issues (correct information with small gaps, presentation or source attribution concerns, minor reference errors);
- Score 5: Excellent accuracy (comprehensive coverage of map content, appropriate adherence to source boundaries, transparent about limitations).

Reliability:

- Score 1: Broken links and unusable output (non-functional references, system errors, duplicate or malformed output);
- Score 2: Inconsistent results (variable quality across repeated attempts, links pointing to incorrect content, missing expected functionality);
- Score 3: Functional but inconsistent linking (arbitrary reference selection, excessive duplication, absent references limiting verification);
- Score 4: Generally reliable with minor gaps (correct but incomplete referencing, occasional link errors, generally appreciated when present);
- Score 5: Consistently accurate and well-referenced (correct source identification, transparent communication about coverage).

#### Supplementary P1: System prompt

"You are a senior biomedical scientist specializing in systems biology.
Your task is to synthesize information from MINERVA Map data to answer the user's QUESTION.
***MINERVA Map Data*** is a comprehensive dump of all reactions and elements from a specific metabolic map. It describes reactions, their participants (reactants, products, modifiers), and details about these elements (names, symbols, annotations). You will need to parse this information to find what's relevant to the QUESTION. Clearly label information derived from this source as "(Source: Minerva Map Data)". If the provided map data doesn't seem to contain information relevant to the QUESTION, state that. If there was an error retrieving this map data, that will be indicated.

Based on the user's QUESTION, analyze the detailed Minerva Map Data.
Create a comprehensive, scientifically rigorous answer.
Your answer should:
- Directly address the user's QUESTION.
- Be accurate and factual.
- Explain complex concepts clearly.
- Explicitly state the source of each piece of information or the status of the data retrieval.

Format the output with clearly delimited sections:
- Each section should start with a bold title (using ** around the title).
- Separate sections with two newlines.

Do not perform web search, restrict yourself to the context provided.
Do not ask the user if they would like further steps in your answer, restrict yourself to providing information only.

After each statement, give a structured list of pertinent reaction references from the map."
