## Supplementary U1 for "User-driven development and evaluation of an agentic framework for analysis of large pathway diagrams"

### Supplementary U1: Llemy default user prompts

Below is the list of pre-constructed example user prompts in Llemy. These prompts were specifically designed to serve as examples that a user could ask the system, having in mind its potential strengths and areas for improvement. By providing standard prompts, we aimed at having a corpus of queries that could give us evaluation results with a smaller variability in the prompts, allowing for a clearer evaluation of the characteristics of the system with a smaller influence of prompt variation. It is important to mention that these were suggestions, and the users were free to ask any prompt, with any structure they wanted.

- What is the scope of this map? Give me a brief summary of the biology represented.
    - With this prompt, we aimed at testing the capacity of global synthesis of Llemy. It was designed to evaluate whether the LLM could condense different map sizes, from hundreds to thousands of nodes and edges, into a coherent narrative.

- What is the overall scope of the disease map (molecular, cellular, tissue, or organism-level)?
    - Similarly, this prompt aimed to evaluate Llemy's capacity for abstraction, however with a slightly different prompt. It also adds multi-level complexity to the expected answer.

- What are the inputs, regulators, and phenotypic outputs of this system?
    - This prompt would need the system to identify the interaction's structure, discriminating inputs, outputs and modifiers. It also tests if the system understands the node annotation in the systems biology standards used in the diagrams (e.g, phenotype nodes), or is able to interpret labels and annotations as a specific entity.

- Which triggers initiate the possible pathological response represented by this map, and which drivers maintain it?
    - This prompt was designed to test Llemy's ability to identify upstream initiators of key pathways leading to a pathological phenotype represented in a map.

- Which regulatory checkpoints limit over-activation of the pathways leading to pathological phenotypes?
    - This prompt was intended to test the system's ability to identify limiting checkpoints, such as bottleneck nodes, which usually are rate limiting for signal propagation within a network. For some maps representing baseline physiological mechanisms, it would also test if the system is extrapolating quantitative interpretations based on qualitative representations.

- How does the microenvironment (inflammatory and metabolic) modulate core pathways?
    - This prompt evaluates contextual sensitivity of map components, and the system's ability to integrate its trained knowledge. Llemy first needs to identify the "core pathways" of the map, possibly reasoning the diagram's contents

against its overall scope. Then evaluate and summarise how the microenvironment, whenever represented in a map, modulates these pathways.

- Are there any inter-organelle interactions (nucleus, mitochondria, membrane, ER) mapped?
    - One of the characteristics of the maps hosted in MINERVA is the use of compartmentalisation, at different granularity, to represent, and sometimes organise, pathway information. Llemy needs to identify these compartments and assess how the processes represented within the organelle compartments interact.

- Which stress pathways (DNA damage, ER stress, oxidative stress, unfolded protein response) are represented in the map?
    - This was designed to assess the model’s recognition and mapping of specific pathways, compartments, and annotations, and also its ability to link map components to its “understanding” of these biological processes.

- Which sentinel nodes serve as proxies for system state?
    - This prompt would require the system to understand the topography of the network, identifying hub nodes in pathways that support the homeostasis of what is represented in the map.

- Does the map capture temporal aspects (e.g. early vs late disease stages)?
    - Many molecular interaction maps are usually static, however, they can depict different stages of a given process (e.g., Atlas of Inflammation Resolution). This prompt was aimed to test the system's ability to identify these differences by interpreting the information retrieved from the map, and/or comparing map data with its trained data.

- Is the mapped system tissue- or cell-type specific? Or are there multiple tissues or cell-types represented?
    - This prompt aimed at identifying Llemy’s ability to interpret and summarise compartment annotation and submap structure.
