## Supplementary U2 for "User-driven development and evaluation of an agentic framework for analysis of large pathway diagrams"

### Supplementary U2: User survey questions

#### Llemy user survey

Please share your experiences when using Llemy, an LLM-based tool for exploration and summarising disease maps. We need your feedback to further develop LLM functionalities.

- please track your time when working with the app, you'll be asked for how much time you've spent, and to estimate your time saved (see below).
- if possible, try to perform a similar task using the map and online resources only, to estimate how much time would it take to reach similar conclusions.

How to:

- Use Llemy (https://llemy.vhp4safety.nl) - you need to provide a valid OpenAI API key.
- Set up your password and use the hash code (see image below) you've obtained for this questionnaire (question 1). Your password will not be stored, make sure you remember it if you use the app again.
- Answer the questions below.

Questions marked with a red asterisk (*) are mandatory.

Where to find the hash code:

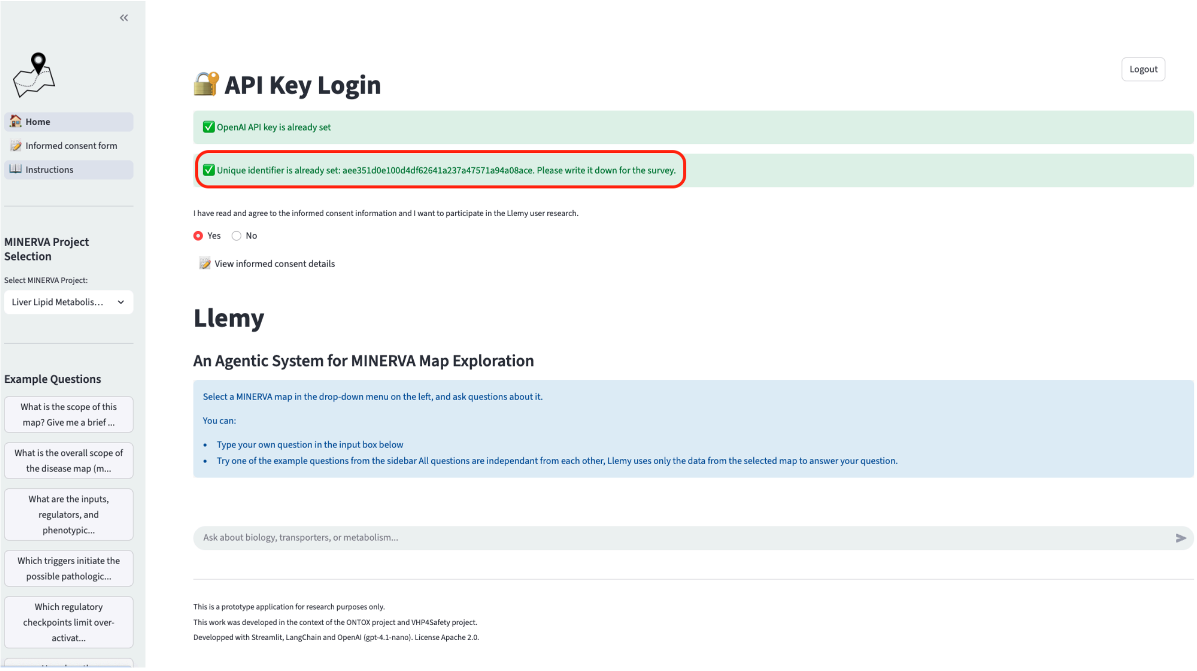


1. Hash identifier provided after login (allows us to relate your answers and your prompts in Llemy): ___________________________________

##### *About you*

Background, experience with LLMs, experience with pathway diagrams

2. What is your experience with pathway diagrams?*

- ☐ None - I never worked with pathway diagrams
- ☐ User - I use pathway diagrams for research
- ☐ Curator - I build pathway diagrams
- ☐ Developer - I develop tools for browsing, building, or analysis of pathway diagrams
- ☐ Other: ________________________________

3. What is your experience with LLMs?*

- ☐ None
- ☐ Basic user - I ask chatGPT for stuff
- ☐ Intermediate user - I spend some time prompting tasks
- ☐ Advanced user - I'm a prompt engineer and have a team of LLM agents working for me
- ☐ Developer - I have trained my own LLM(s)
- ☐ Other: ________________________________

#### *Working with the app*

Your experience working with Llemy.

4. For how long did you work with the app?*

- ☐ 15 min or less
- ☐ Between 15 and 30min
- ☐ Between 30min and an hour
- ☐ More than an hour

5. What tasks did you perform?*

- ☐ Summarise entire diagrams
- ☐ Summarise mechanisms involving specific elements or interactions
- ☐ Identify mechanisms of interest (e.g. downstream targets, signalling pathways, etc.)
- ☐ Identify knowledge gaps
- ☐ Identify new elements interacting with the contents of the map (e.g. ligands, enzymes, transcription factors, protein interactors)
- ☐ Other: ________________________________

6. How easy was it to use the app?*

Very difficult

- ☐ 1
- ☐ 2
- ☐ 3
- ☐ 4
- ☐ 5

Very easy

7. Comments on ease of use (if any): ________________________________

#### *Productivity, reliability and usefulness*

Estimate the time and effort saved when performing the tasks.

8. How much time did you save? (try to estimate by comparing with a similar task you would do yourself)*

- ☐ Time lost (reading and verifying output took more time than you would spend on the task)
- ☐ No time gain (reading and verifying output took as much time as you would spend on the task)
- ☐ Up to 50% time gain (every hour of work saves 30 min of your time)
- ☐ Up to 100% time gain (every hour of work saves an hour of your time)
- ☐ Up to 200% time gain (every 30 min of work saves an hour of your time)
- ☐ More than 200% time gain
- ☐ Other: ________________________________

9. Comments on time saved (if any): ________________________________

10. How much does the structure of the output change between similar prompts?

Significant change between queries

- ☐ 1
- ☐ 2
- ☐ 3
- ☐ 4
- ☐ 5

Completely reproducible results

11. Comments on output change between prompts (if any): ________________________

12. What is your overall assessment of the usefulness of the app?*

Not useful

- ☐ 1
- ☐ 2
- ☐ 3
- ☐ 4
- ☐ 5

Very useful

13. Comments on overall assessment (if any): ________________________________

#### *Final comments*

14. Suggestions for improvement (interface, output, etc): ___________________________
