## Supplementary U3 for "User-driven development and evaluation of an agentic framework for analysis of large pathway diagrams"

### Supplementary U3: User evaluation metrics

| ID | Category | Response time (seconds) | Feedback accuracy score | Feedback conciseness score | Feedback reliability score |
|---|---|---|---|---|---|
| 001 | Analyse | 28.2163 | 5 | 3 | 4 |
| 002 | Summarise | 22.4099 | 2 | 1 | 2 |
| 003 | Find | 78.1817 | 5 | 4 | 5 |
| 004 | Find | 222.0802 | 4 | 2 | 5 |
| 005 | Find | 412.6278 | 2 | 3 | 2 |
| 006 | Summarise | 38.4972 | 5 | 4 | 5 |
| 007 | Analyse | 33.1146 | 5 | 4 | 3 |
| 008 | Summarise | 42.2872 | 3 | 3 | 3 |
| 009 | Analyse | 43.8341 | 4 | 3 | 3 |
| 010 | Find | 81.2427 | 3 | 3 | 3 |
| 011 | Analyse | 21.3286 | 5 | 1 | 5 |
| 012 | Analyse | 21.3286 | 5 | 2 | 5 |
| 013 | Summarise | 42.8327 | 3 | 3 | 3 |
| 014 | Summarise | 225.4019 | 1 | 3 | 3 |
| 015 | Find | 35.3072 | 4 | 3 | 4 |
| 016 | Find | 456.1268 | 1 | 1 | 1 |
| 017 | Out of Scope | 27.4588 | 4 | 3 | 5 |
| 018 | Summarise | 82.804 | 5 | 5 | 5 |
| 019 | Summarise | 21.235 | 1 | 3 | 1 |
| 020 | Summarise | 20.9294 | 2 | 3 | 2 |
| 021 | Summarise | 18.2458 | 3 | 3 | 3 |
| 022 | Find | 244.8034 | 3 | 1 | 1 |
| 023 | Summarise | 21.4183 | 5 | 1 | 5 |
| 024 | Summarise | 25.7209 | 5 | 4 | 4 |
| 025 | Summarise | 76.9444 | 5 | 3 | 5 |
| 026 | Summarise | 34.0399 | 4 | 4 | 4 |
| 027 | Analyse | 76.4653 | 1 | 2 | 2 |
| 028 | Find | 81.9757 | 5 | 5 | 5 |
| 029 | Analyse | 82.5072 | 4 | 2 | 1 |
| 030 | Find | 264.0319 | 4 | 2 | 4 |
| 031 | Analyse | 17.968 | 5 | 3 | 5 |
| 032 | Find | 18.5621 | 1 | 3 | 1 |

| | | | | | |
|---|---|---|---|---|---|
| 033 | Analyse | 28.3291 | 4 | 4 | 4 |
| 034 | Summarise | 35.8252 | 2 | 3 | 3 |
| 035 | Analyse | 151.7659 | 1 | 1 | 3 |
| 036 | Analyse | 237.554 | 1 | 1 | 3 |
| 037 | Summarise | 462.1365 | 1 | 1 | 1 |
| 038 | Summarise | 36.3324 | 4 | 3 | 4 |
| 039 | Analyse | 14.8958 | 4 | 4 | 4 |
| 040 | Analyse | 86.5625 | 2 | 4 | 2 |
| 041 | Analyse | 44.2533 | 5 | 4 | 5 |
| 042 | Analyse | 37.787 | 3 | 3 | 3 |
| 043 | Analyse | 78.5608 | 3 | 4 | 4 |
| 044 | Analyse | 74.8395 | 2 | 3 | 3 |
| 045 | Find | 44.6514 | 3 | 3 | 3 |
| 046 | Analyse | 20.4002 | 4 | 4 | 4 |
| 047 | Analyse | 77.1516 | 2 | 2 | 3 |
| 048 | Analyse | 33.0616 | 4 | 4 | 4 |
| 049 | Summarise | 36.3417 | 3 | 3 | 4 |
| 050 | Find | 26.4415 | 3 | 3 | 3 |
| 051 | Find | 30.4079 | 4 | 4 | 4 |
| 052 | Summarise | 30.0975 | 5 | 5 | 5 |
| 053 | Analyse | 19.6508 | 3 | 3 | 3 |
| 054 | Summarise | 84.2385 | 4 | 4 | 4 |
| 055 | Summarise | 78.545 | 2 | 2 | 4 |
| 056 | Summarise | 21.7004 | 5 | 1 | 5 |
| 057 | Summarise | 25.5644 | 5 | 1 | 5 |
| 058 | Summarise | 30.3399 | 4 | 4 | 4 |
| 059 | Summarise | 34.7802 | 5 | 4 | 5 |
| 060 | Find | 34.3063 | 3 | 4 | 4 |
| 061 | Summarise | 437.6224 | 1 | 2 | 1 |
| 062 | Find | 20.5333 | 3 | 1 | 3 |
| 063 | Summarise | 36.8527 | 2 | 3 | 2 |
| 064 | Find | 26.9604 | 4 | 3 | 1 |
| 065 | Find | 32.0679 | 3 | 3 | 2 |
| 066 | Find | 18.1711 | 4 | 3 | 3 |
| 067 | Find | 276.6109 | 1 | 1 | 1 |
| 068 | Analyse | 28.4058 | 5 | 4 | 3 |

| | | | | | |
|---|---|---|---|---|---|
| 069 | Analyse | 10.3914 | 1 | 1 | 3 |
| 070 | Find | 929.1113 | 4 | 4 | 4 |
| 071 | Find | 251.653 | 3 | 4 | 4 |
| 072 | Analyse | 28.3122 | 4 | 4 | 3 |
| 073 | Summarise | 31.1276 | 2 | 3 | 2 |
| 074 | Find | 44.8807 | 3 | 4 | 4 |
| 075 | Summarise | 23.1607 | 4 | 5 | 5 |
| 076 | Find | 35.0634 | 5 | 5 | 2 |
| 077 | Analyse | 26.3004 | 4 | 3 | 4 |
| 078 | Analyse | 35.4496 | 5 | 5 | 5 |
| 079 | Analyse | 39.4214 | 4 | 4 | 3 |
| 080 | Analyse | 29.2793 | 3 | 3 | 3 |
| 081 | Summarise | 239.7815 | 2 | 2 | 2 |
| 082 | Find | 27.2643 | 5 | 5 | 4 |
| 083 | Find | 3.7125 | 4 | 4 | 4 |
| 084 | Summarise | 36.7102 | 4 | 4 | 4 |
| 085 | Analyse | 6.1923 | 5 | 5 | 5 |
| 086 | Find | 191.712 | 4 | 2 | 4 |
| 087 | Find | 191.712 | 4 | 1 | 4 |
| 088 | Analyse | 20.777 | 5 | 5 | 4 |
| 089 | Analyse | 31.2476 | 5 | 5 | 5 |
| 090 | Analyse | 19.177 | 4 | 5 | 5 |
| 091 | Summarise | 34.0604 | 3 | 3 | 2 |
| 092 | Summarise | 228.1139 | 3 | 3 | 3 |
| 093 | Analyse | 232.7486 | 4 | 1 | 3 |
| 094 | Summarise | 37.6147 | 5 | 5 | 4 |
| 095 | Summarise | 37.6147 | 5 | 5 | 5 |
| 096 | Summarise | 34.4011 | 4 | 3 | 4 |
| 097 | Summarise | 36.9448 | 5 | 5 | 4 |
| 098 | Summarise | 32.5815 | 2 | 3 | 3 |
| 099 | Summarise | 35.5714 | 3 | 4 | 3 |
| 100 | Find | 35.7028 | 4 | 4 | 2 |
| 101 | Find | 34.8528 | 4 | 4 | 4 |
| 102 | Find | 34.3248 | 3 | 2 | 3 |
| 103 | Find | 23.3839 | 5 | 4 | 5 |
| 104 | Find | 28.2461 | 3 | 3 | 3 |

| | | | | | |
|---|---|---|---|---|---|
| 105 | Find | 13.7126 | 1 | 2 | 3 |
| 106 | Find | 22.5613 | 3 | 3 | 3 |
| 107 | Find | 20.2563 | 3 | 3 | 4 |
| 108 | Analyse | 14.3613 | 1 | 4 | 1 |
| 109 | Find | 19.7944 | 5 | 4 | 3 |
| 110 | Summarise | 76.0591 | 3 | 3 | 2 |
| 111 | Summarise | 25.227 | 4 | 4 | 4 |
| 112 | Summarise | 23.2739 | 3 | 3 | 3 |
| 113 | Summarise | 8.0092 | 5 | 4 | 5 |
| 114 | Summarise | 79.6202 | 2 | 3 | 3 |
| 115 | Summarise | 16.0143 | 3 | 3 | 3 |
| 116 | Summarise | 86.6634 | 5 | 5 | 5 |
| 117 | Summarise | 42.7544 | 4 | 5 | 3 |
| 118 | Summarise | 44.859 | 4 | 4 | 4 |
| 119 | Summarise | 21.3974 | 5 | 3 | 5 |
| 120 | Summarise | 33.542 | 4 | 4 | 4 |
| 121 | Summarise | 36.1631 | 4 | 3 | 4 |
| 122 | Summarise | 23.4826 | 1 | 2 | 3 |
| 123 | Summarise | 28.9572 | 4 | 3 | 3 |
| 124 | Analyse | 6.3152 | 4 | 4 | 3 |
| 125 | Find | 28.7738 | 5 | 4 | 5 |
| 126 | Summarise | 14.9326 | 2 | 2 | 3 |
| 127 | Analyse | 33.975 | 5 | 3 | 3 |
| 128 | Analyse | 90.2133 | 5 | 4 | 4 |
| 129 | Analyse | 40.8464 | 3 | 3 | 3 |
| 130 | Summarise | 34.2553 | 4 | 3 | 4 |
| 131 | Analyse | 225.6587 | 3 | 4 | 4 |
| 132 | Analyse | 156.689 | 3 | 1 | 3 |
| 133 | Analyse | 83.6239 | 3 | 4 | 3 |
| 134 | Find | 26.9125 | 4 | 3 | 4 |
| 135 | Find | 75.1455 | 4 | 3 | 4 |
| 136 | Find | 83.9212 | 4 | 4 | 2 |
| 137 | Analyse | 44.1013 | 4 | 3 | 3 |
| 138 | Find | 252.1148 | 1 | 1 | 1 |
| 139 | Analyse | 84.3028 | 5 | 3 | 2 |
| 140 | Summarise | 27.4738 | 4 | 4 | 4 |

| | | | | | |
|---|---|---|---|---|---|
| 141 | Analyse | 242.6845 | 1 | 1 | 1 |
| 142 | Find | 20.1116 | 3 | 3 | 2 |
| 143 | Analyse | 9.3164 | 4 | 4 | 4 |
| 144 | Analyse | 50.6857 | 5 | 4 | 5 |
| 145 | Analyse | 37.2194 | 3 | 4 | 3 |
| 146 | Analyse | 39.5466 | 4 | 4 | 3 |
| 147 | Analyse | 43.6011 | 5 | 4 | 5 |
| 148 | Find | 36.5658 | 4 | 4 | 5 |
| 149 | Find | 265.6204 | 1 | 1 | 1 |
| 150 | Find | 27.5797 | 4 | 4 | 4 |
| 151 | Find | 42.6832 | 5 | 4 | 5 |
| 152 | Find | 166.7365 | 4 | 4 | 3 |
| 153 | Find | 24.0201 | 3 | 4 | 4 |
| 154 | Analyse | 49.0015 | 5 | 3 | 5 |
| 155 | Analyse | 36.4246 | 2 | 3 | 3 |
| 156 | Analyse | 27.747 | 3 | 3 | 3 |
| 157 | Summarise | 26.991 | 5 | 5 | 5 |
